# *Is* EEG is better left alone?

**DOI:** 10.1101/2023.06.19.545602

**Authors:** Alain de Cheveigné

**Author notes:** Author’s contribution statement The author, Alain de Cheveigné, is responsible for all aspects of this paper.

## Abstract

A recent article (Delorme, A., 2023, EEG is better left alone. *Scientific Reports*, **13**, 2372. https://doi.org/10.1038/s41598-023-27528-0) proposed a metric to determine the benefit of applying pre-processing methods to EEG data. Using that metric, it concluded that most pre-processing methods do *not* improve data quality. The question is revisited in this short paper, using a sample of the same data as used by that paper. It is argued that the metric is of limited applicability, and that in some situations pre-processing might be critical to make good use of the data.

Note: This paper was submitted to Scientific Reports as a commentary to the paper cited and rejected without review.

## Introduction

Electro-encephalography (EEG) offers a window into brain activity that is often clouded by noise and artifacts. To improve their quality, EEG data may be *pre-processed* as a first step in their analysis, but there are many methods available, each with multiple parameters, and it is not always clear which (if any) actually improves the outcome. Adjusting the pre-processing chain, based on how well it deals with a dataset, introduces unwanted “ researcher degrees of freedom” that may invalidate statistical tests [1] (Simmons et al 2021), on top of being unwieldy for datasets with hundreds of subjects. A metric that can be applied across the board, to any dataset or pre-processing chain to help decide what to include in the pipeline, is useful. [2] Delorme (2023) proposes such a metric. On its basis, he draws a sobering conclusion: most pre-processing methods do *not* improve data quality, hence his title: “ *EEG is better left alone*”. This commentary questions the generality of the metric and the conclusion.

The metric involves counting the *number of significant channels* for a statistical test applied to an effect (for example a difference between two conditions during some time window). The rationale is appealing: thanks to the current spread, every electrode should pick up a signal from the brain source that underlies the effect, with a signal-to-noise ratio (SNR) that reflects the level of noise at that electrode. A better SNR should increase the number of channels for which the statistical test is positive, resulting in a higher score. At first glance it seems a straightforward, no-nonsense way to quantify the effectiveness of pre-processing. However, it raises several issues.

First, information is reduced to a single bit for each channel, and the resulting N bits are further reduced to log_2_N bits by the counting (Fig. 1 a). Quantization adds variability (“quantization noise”) that squanders the statistical power of a second-level statistical test to determine if a difference in metric is significant or not, or higher-level tests to assess generality across subjects or datasets. In addition, the measure might be brittle: a small change in SNR near the threshold might induce a large change in metric, while a large change far from the threshold might leave it unchanged.

**Fig. 1.**
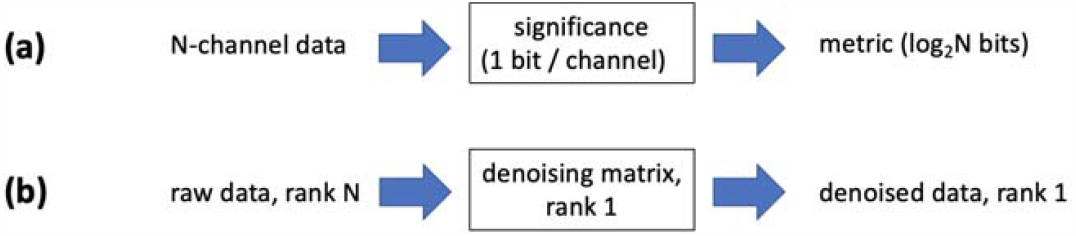
Two issues with the significant-channel-count metric. (a) The significance test reduces information to one bit per channel, further reduced to log_2_N bits by counting (e.g. 6 bits for 64 channels), so the metric is roughly quantized. (b) Multiplying the EEG data matrix by a denoising matrix of rank one yields a denoised matrix of rank one: all channels are the same to a scaling factor. The metric then can take only two values: 0 or 100 %.

Second, the metric runs into difficulty for a pre-processing algorithm that reduces the rank of the data to one (Fig. 1b). For example, Independent Component Analysis (ICA, [3] Makeig et al 1996, [4] Delorme et al. 2012) might capture within a single component the brain source responsible for the effect. The best strategy then is to project that component back to sensor space to obtain “clean” data, in which case *all the channels carry the same signal* to a multiplicative factor. The metric then can take only one of two values (0 or 100 %), again making it a crude measure of quality when applied in this situation.

Third, the metric is contingent on an effect. It is not clear whether it should (or can) be applied afresh to each new effect or whether conclusions drawn from one effect can be generalized across the board. Delorme’s paper assumes the latter, as it draws general conclusions about pre-processing algorithms and suites on the basis of three effects observed each within a different dataset. For two datasets, brain activity was most likely dominated by motor activity (subjects responded to one condition and not the other), which is usually strong and in little need of pre-processing, so those conclusions might not generalize to more subtle brain activity.

## Methods

To illustrate this point, the data of one subject of the third dataset (“auditory oddball”, doi:10.18112/openneuro.ds003061.v1.1.2) are analyzed here. The process is briefly sketched, the important messages being that pre-processing can be effective, that it may involve multiple steps, and that choosing and tuning these steps may involve the investigator’s judgment. In other words, analysis is not always amenable to a standard pipeline or easily evaluated by a blanket metric.

In the recording sessions, listeners heard a succession of tones at one frequency (1 kHz, the “Standard”) interspersed with less frequent tones at another frequency (500 Hz, the “Oddball”) to which they had to respond by a button press (see [2] Delorme 2023 for further details). The effect used by the metric is a difference in brain response between Standard and Oddball. An Oddball stimulus might indeed elicit a “mismatched negativity” (MMN, [5] Näätänen et al. 2007), here likely compounded by motor activity.

The analysis proceeds in steps. After an initial inspection, the data (time X channels) are divided into trial matrices (time X channels X trials), one per event type (Standard onset, Oddball onset, button press), each trial spanning the interval [-2s, +2s] relative to the event. Next, data are detrended by fitting a linear function to each channel of each trial that is then subtracted [6] (de Cheveigné and Arzounian, 2018), after which each trial is truncated to a shorter interval, e.g. [-1s, +1s]. This is to suppress slow trends, ubiquitous in EEG, due to polarization phenomena at the skin/electrolyte/electrode interface (Fig. 2 a). Next, power line artifacts are removed using the ZapLine algorithm [7] (de Cheveigné 2020) (Fig. 2b). Interestingly, the line frequency was found to range between 49 and 51 Hz (EEG were recorded in a location with unstable line noise), so default parameters required adjustment. The data are then re-referenced to the mean, and the pre-event mean is removed.

**Fig. 2.**
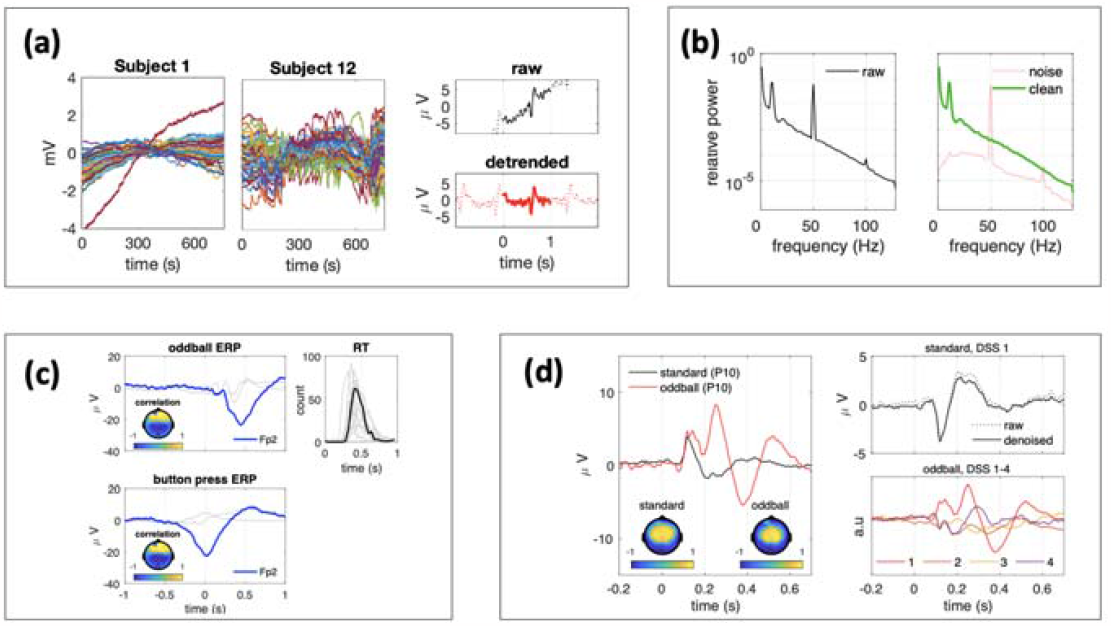
Stages in pre-processing. (a) A major source of artifacts in EEG is a slow trend, which can be variously suppressed by high-pass filtering or detrending. A scheme applicable to one set of data (e.g. Subject 1, smooth trends) may be inadequate for another (e.g. Subject 12, erratic trends). Here, a linear trend is removed from each trial over an extended epoch, to avoid affecting the ERP shape. (b) Power line artifacts are removed using the Zapline algorithm. (c) ERPs time-locked to the Oddball stimulus are suspiciously similar (amplitude, topography) to those time-locked to the Button press, suggesting that both may be dominated by motor activity, consistent with the histogram of reaction times (right). Topographies represent the correlation between the trial-averaged component and each trial-averaged electrode signal. Motor activity may be attenuated by projecting out the three components most reliably time-locked to the button press (2nd and 3rd are shown in light gray), as a form of “denoising”. (d) DSS is applied separately to denoised Standard and Oddball trial matrices, and the most reliable components of each are aligned using CCA. The first CCs (best aligned) for the Standard and Oddball are compared in the left panel. The similarity of topographies and initial time course suggests that the same brain sources are involved, but with a time course that diverges beyond ∼200 ms. These responses are unlikely to have been distorted by denoising (upper right plot). Additional stimulus-locked components are specific to the Oddball response (lower right plot). Together with the button-press-locked components of panel c, they span the space of sensory, decisional, and motor activity within the brain that is deployed when responding to an Oddball stimulus. Pre-processing helps reveal this activity.

Next, Denoising Source Separation (DSS, [8] Särelä and Valpola 2005, de Cheveigné and Parra 2014) is applied to each trial matrix, separately, to isolate components time-locked to the event (Standard, Oddball, button press). Stimulus-locked components are found to be very different between Standard and Oddball but similar between Oddball and button press (Fig. 2c), suggesting that event-related potentials (ERP) for Oddball are mainly due to brain activity related to the temporally adjacent button press (see histogram of reaction times in Fig. 2c), which prevents interpretation in terms of a sensory response specific to rare stimuli. The rest of the analysis attempts to circumvent this obstacle.

Trial matrices for both stimuli are denoised by projecting out the first three components from the previous DSS analysis. The aim is to suppress the button-press-locked components and reveal sensory features specific to Oddball (if any). This denoising is applied to both Oddball (for which it is necessary) and Standard (for which it is not) for consistency. DSS is then applied anew to each denoised stimulus-locked trial matrix to isolate components reliably time-locked to each of these two stimuli. A subset of reliable components is selected for each (3 from Standard, 5 from Oddball), and these subsets are finally aligned using Canonical Component Analysis (CCA) to facilitate comparison. The first canonical components (CCs) are shown in Fig. 2 (d, left).

The responses, initially quite similar, diverge radically from ∼200 ms onwards, with a topography (plotted here as the correlation coefficient between this component and each EEG channel, both trial-averaged, calculated over the entire epoch) similar to an auditory MMN [10] (e.g. Garrido et al. 2007). The Oddball response includes additional components (Fig. 2d lower right) that either differ in form between Oddball and Standard, or exist only within the Oddball response. Together with the button-press-locked components, they span a rich continuum of brain activity that presumably reflects a chain of sensory processing, decision, motor preparation, and execution, from Oddball detection to button press. The purpose here is neither to investigate this activity, nor to defend the choice of analysis or parameters, nor to address the many methodological questions that the analysis raises or the controls that it calls for. Instead, I wish to make some more general points.

The ambition of brain data analysis should be to push the limits of what can be observed in complex brain activity. Pre-processing should be measured on the scale of this ambition. The distinction between pre-processing and data analysis is not clear-cut. Here, each step revealed something about the data (e.g. presence of drift, power-line artifact, or motor response), and also cleared the way for the next step. The final DSS would have failed without the initial DSS, which would have failed without detrending. Each step addresses a different artifact, and is helpful only if that artifact is present (and disruptive), and thus it should not come as a surprise if including a particular method in the pipeline does not always improve the outcome.

Pre-processing involves a chain of decisions, each with proximal and distant consequences (as in a game of chess). High-pass filtering might effectively suppress the slow drift, but distort the ERPs [11] (de Cheveigné and Nelken 2019), hence the decision here to use trial-based detrending instead. Re-referencing does not improve the outcome of component analysis (on the contrary, it reduces the rank of the data by one), but it makes topographies more informative. Baseline correction (removal of pre-event mean) is justified on logical grounds. Here, some of these decisions were made “on the fly” as the analysis progressed. In other words, they were not all planned in advance, and it is not clear that they could (or should) be. For example, a pre-planned analysis contrasting Standard and Oddball ERPs (as implicit in [2] Delorme 2023) might fail to notice that the latter are dominated by motor activity, and miss the opportunity to analyze them further.

There is a fundamental tension between an efficient, easy-to-report, hands-off, pre-planned (and pre-registered) analysis pipeline and a data-aware analysis process that adapts to artifacts discovered in the data (and confounds discovered in the paradigm), at the cost of introducing “researcher degrees of freedom”. Both are eminently desirable but not entirely compatible, a case where you cannot “have your cake and eat it too”. It has been suggested to brand the former “confirmatory” and the latter “exploratory” [12] (Nosek et a 2018), but this may not fully resolve the tension [13] (de Cheveigné, 2023).

## Summary

Delorme’s (2023) study addressed a real need. To analyze brain data in a principled manner, and avoid tinkering and unwanted degrees of freedom, it is useful to develop standardized pipelines and metrics to evaluate them. The metric proposed by Delorme (2023) is an important step in this direction.

A more general question is whether the goal of fully “hands-off” data analysis is feasible. While the goal is worth approximating, two arguments can be put forward to temper optimism. One is that no amount of testing can guarantee correctness, a point famously made by Edsger Dijkstra (1970) in the context of software [14]. The second is that a feature of scientific research is that it ventures into unchartered territory where precedent is sparse. While standardization makes good sense in a well-trodden field, it is possibly less useful at the cutting edge.

The answer to the question in the title is: “it depends”. Each pre-processing tool targets an artifact, and if that artifact does not occur within the data, then that tool is best not applied. In some situations, however, well-designed pre-processing steps can help make sense of the recordings. In those situations, EEG is best *not* left alone.

## Acknowledgments

This work was funded in part by FrontCog grant ANR-17-EURE-0017. Arnaud Delorme made useful comments on a previous version.

## Code availability

A script to reproduce Fig. 2 (and document details of the analysis) is available from audition.ens.fr/adc/NoiseTools/src/NoiseTools/EXAMPLES/ScientificReports_2023/ based on routines from the NoiseTools toolbox, audition.ens.fr/adc/NoiseTools/.

## References

[1] Simmons, J., Nelson, L., Simonsohn, U. False-Positive Psychology: Undisclosed Flexibility in Data Collection and Analysis Allows Presenting Anything as Significant. Psychological Science, 22, 1359–1366 (2021).

[2] Delorme, A. EEG is better left alone. Scientific Reports, 13, 2372. https://doi.org/10.1038/s41598-023-27528-0 (2023)

[3] Makeig, S. et al. (1996) Independent component analysis of electroencephalographic data. Adv. Neural Inf. Process. Syst. 8, 145–151

[4] Delorme, A., Palmer, J., Onton, J., Oostenveld, R., & Makeig, S. Independent EEG Sources Are Dipolar. PLoS ONE, 7, e30135. https://doi.org/10.1371/journal.pone.0030135 (2012).

[5] Näätänen R, Paavilainen P, Rinne T, Alho K. The mismatch negativity (MMN) in basic research of central auditory processing: a review. Clin. Neurophysiol., 118, 2544–90 (2007).

[6] de Cheveigné, A., Arzounian, D. Robust detrending, rereferencing, outlier detection, and inpainting for multichannel data. NeuroImage, 172, 903–912 10.1016/j.neuroimage.2018.01.035 (2018).

[7] de Cheveigné, A. ZapLine: A simple and effective method to remove power line artifacts, Neuroimage 207, 116356, doi: 10.1016/j.neuroimage.2019.116356 (2020).

[8] Särelä, J., Valpola, H., Denoising source separation. J. Mach. Learn. Res. 6, 233–272 (2005).

[9] de Cheveigné, A., Parra, L. Joint decorrelation: a versatile tool for multichannel data analysis. NeuroImage, 98, 487–505, doi: 10.1016/j.neuroimage.2014.05.068 (2014).

[10] Garrido, M. I., Kilner, J. M., Stephan, K. E., & Friston, K. J. The mismatch negativity: A review of underlying mechanisms. Clinical Neurophysiology, 120, 453–463. https://doi.org/10.1016/j.clinph.2008.11.029 (2009).

[11] de Cheveigné, A., Nelken, I. Filters: why, when and how (not) to use them, Neuron 102, 280–293, doi: 10.1016/j.neuron.2019.02.039 (2019).

[12] Nosek, B. A., Ebersole, C. R., DeHaven, A. C., & Mellor, D. T. The preregistration revolution. Proceedings of the National Academy of Sciences, 115 (11),2600–2606. doi: 10.1073/pnas.1708274114 (2018).

[13] de Cheveigné, A. Preregistration: the good, the bad, and the confusing.PsyArXiv https://psyarxiv.com/bcd9t/ (2023).

[14] Dijkstra, E.W. Notes on Structured Programming. T.H.-Report 70–WSK–03, Second edition (available at http://www.cs.utexas.edu/users/EWD/ewd02xx/EWD249.PDF) (1970).

